# scAgeClock: a single-cell transcriptome based human aging clock model using gated multi-head attention neural networks

**DOI:** 10.1101/2025.08.29.673183

**Authors:** Gangcai Xie

**Affiliations:** Institute of Reproductive Medicine, Medical School, Nantong University, Nantong, China; Jiangsu Province Key Laboratory in University for Inflammation and Molecular Drug Target, Nantong University, Nantong, China

**Author notes:** Correspondence: Gangcai Xie, Nantong University, Qixiu Road 19, Nantong, China, 226001.

## Abstract

Aging Clock models have emerged as a crucial tool for measuring biological age, with significant implications for anti-aging interventions and disease risk assessment. However, human aging clock models that offer single-cell resolution and account for cell and tissue heterogeneities remain underdeveloped. This study introduces scAgeClock, a novel gated multi-head attention (GMA) neural network-based single-cell aging clock model. Leveraging a large-scale dataset of over 16 million single-cell transcriptome profiles from more than 40 human tissues and 400 cell types, scAgeClock demonstrates improved age prediction accuracy compared to baseline methods. Notably, the mean absolute error for the best-performing cell type is remarkably low at 2 years. Feature importance analysis reveals enrichment of aging clock genes related to ribosome, translation, defense response, viral life cycle, programmed cell death, and COVID-19 disease. A novel metric, the Aging Deviation Index (ADI) proposed by this study, revealed deceleration of ages in cells with higher differentiation potencies and tumor cells in higher phases or under metastasis, while acceleration of ages was observed in skin cells. Furthermore, scAgeClock is publicly available to facilitate future research and potential implementations.

## INTRODUCTION

Aging is a multifaceted process that encompasses not only visible manifestations, such as graying hair, brittle bones, and wrinkled skin, but also subtle microscopic changes, including telomere shortening, epigenetic dysfunction, genomic instability, and alterations in protein and gene expression[1]. Moreover, aging is a significant risk factor for various human diseases, including Parkinson’s disease, Alzheimer’s disease, heart failure, Type 2 diabetes, and osteoarthritis[2]. Notably, individuals with the same chronological age can exhibit vastly different biological ages, highlighting the importance of precise measurement. Accurately determining an individual’s biological age could confer numerous benefits, including evaluating the efficacy of anti-aging interventions, informing personalized health risk assessments, enabling early detection of aging-related diseases, and motivating healthier lifestyle choices.

Aging clocks are a crucial tool for measuring biological age, and can be developed using various approaches, including macroscopic features such as 3D facial images, eye corner images, brain MRI scans, and psychosocial questionaries, as well as molecular changes like DNA methylation, gene expression, and metabolic shifts[3]. The first molecular-level aging clock based on genome-wide large-scale cohorts was the DNA methylation-based model introduced by Hannum et al.[4] and Horvath[5] over a decade ago. Since then, numerous omics datasets have been used for aging clock development, including epigenomics, transcriptomics, proteomics, metabolomics, microbiomics, and glycomics[6]. However, most early methods relied on bulk-level samples for omics measurement, neglecting the importance of cell-type heterogeneity and changes in cell-type composition during the aging process. A recent meta-analysis highlighted this limitation, revealing that the same aging clock method can yield different conclusions about age-acceleration in certain diseases when cell-type heterogeneity is either considered or ignored[7].

The advent of single-cell sequencing technologies has led to the development of several aging clock methods based on single-cell omics datasets. Some studies have focused on building single-cell aging clock models in mice. For instance, Buckley et al. constructed a cell-type-specific brain aging clock using single-cell RNA sequencing (scRNA-seq) data from 28 mice spanning the adult mouse lifespan (3-29 months), demonstrating its potential to quantify transcriptomic rejuvenation[8]. Subsequent work utilizing the Tabula Muris Senis datasets led to the development of SCALE, a statistical method for single-cell aging clock estimation that integrates both cell-type and tissue information for biological age prediction[9]. Moreover, Bonder et al. developed a single-cell DNA methylation aging clock model based on peripheral blood samples, highlighting epigenetic heterogeneity during aging process[10].

Other recent studies have focused on developing human single-cell aging clock models based on scRNA-seq datasets, with a primary emphasis on utilizing human blood samples. For example, Zhu et al. developed a human PBMC scRNA-seq-based aging clock model using datasets from 17 natural aging cohorts and 7 supercentenarians [11]. Notably, this model relied on cell type proportion information rather than single-cell gene expression for its construction and revealed the significant contribution of ribosome genes to the aging process. In a subsequent study, a larger scRNA-seq dataset comprising over 1 million blood cells from 508 human donors was utilized to train an elastic net linear regression model [12]. Although this approach employed single-cell gene expression data for biological age prediction, the best-performing cell type achieved only moderate accuracy, with a mean absolute error (MAE) ranging from 8 to 10 and a Pearson correlation coefficient of approximately 0.5.

Previous epigenetic aging clock models have primarily employed regularized linear regression[13], while aforementioned single-cell transcriptomic aging clocks have been built using partial least squares regression[11] or elastic net linear regression[12]. Inspired by the successful applications of neural network models in other fields, deep neural networks have also been adapted for aging clock construction. Examples include the inflammatory aging clock iAge[14], which leverages immune phenotyping datasets; ThermoFace[15], a deep learning-based aging clock model that utilizes thermal facial images; sc-ImmuAging[16], an immune aging clock model based on PBMC scRNA-seq data and built using a deep learning framework (PointNet[17]). However, current human aging clock models are predominantly focused on data from blood tissue, highlighting the need for more comprehensive models that incorporate data from multiple human organs and tissues.

The past few years have seen a significant accumulation of single-cell RNA sequencing datasets. For instance, the public database CZ CELLxGENE Discover[18] provides access to over 70 million single-cell transcriptomic profiles from human samples, spanning ages from less than 1 year old to around 100 years old. Meanwhile, large language models (LLMs) have made remarkable progress in the past two years, achieving human-level performance on various tasks. Notably, most of these LLMs employ Transformer-based neural network architectures[19], as seen in examples such as ChatGPT-4[20] and DeepSeek-V3[21]. Transformer architecture relies heavily on neural networks with attention mechanisms, and multi-head attention neural networks are a crucial component of this design[19].

In this work, I leverage the large-scale human scRNA-seq datasets collected by CZ CELLxGENE Discover to propose a novel aging clock model, scAgeClock, based on gated multi-head attention neural networks. ScAgeClock demonstrates higher accuracy compared to baseline methods and enables single-cell level transcriptomic biological age prediction for major human tissues and cell types, thereby expanding its applications to various scenarios in aging research. Moreover, I introduce a new metric index, the Aging Deviation Index (ADI), which can capture aging-related cell dynamics in cells from both normal and disease tissues based on single-cell-level biological age predictions generated by scAgeClock. Overall, the scAgeClock presented in this study not only serve as a valuable tool for single-cell level biological age prediction but also provides new insights into disease-related cell dynamic changes.

## RESULTS

### Overview of the scAgeClock model and Dataset Composition

In this study, I developed a novel aging clock model, scAgeClock, based on single-cell RNA sequencing (scRNA-seq) data. The model leverages a gated multi-head attention neural network architecture consisting of five layers: embedding, feature gating, feature projection, multi-head attention, and a fully connected neural network layer with age prediction output (Figure 1A). The input data for training the model consisted of normal-tissue human scRNA-seq data from CZ CELLxGENE after filtering by multiple criteria. Specifically, the total number of cells in CZ CELLxGENE for human species is 74,322,510. After selecting cells from normal tissues and primary cohorts, 30,197,419 cells remained. The data was further filtered by age information, resulting in 16,497,049 cells with accurate age information. Subsequent filtering steps were applied to assay type (resulting in 16,496,599), cell type (16,491,591 cells) and donors (16,231,138 cells).

**Figure 1.**
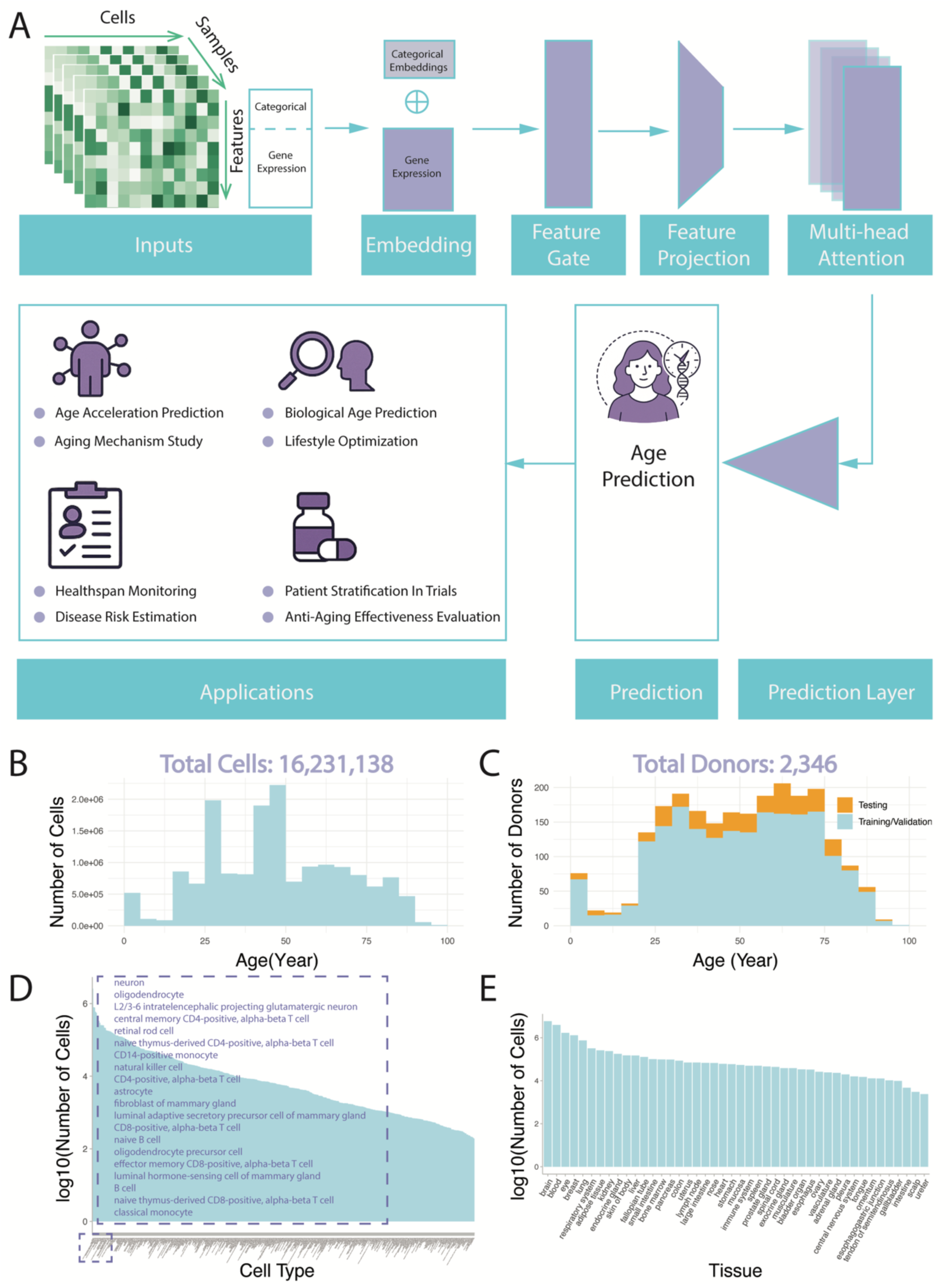
Schematic overview of scAgeClock neural network architecture and dataset characteristics. (A) Illustration of the Gated Multi-head Attention (GMA) neural network architecture utilized in scAgeClock, highlighting its proposed applications in various fields. (B) Age distribution of cells filtered from the CZ CELLxGENE normal human single-cell RNA sequencing (scRNA-seq) dataset. This dataset is used for model training, validation and testing. (C) Donor age distribution for each dataset used in model development. (D-E) Cell-type-level (D) or tissue-level (E) cell-count distribution (log10-transformed) for the datasets in model development.

The final filtered dataset encompassed a diverse range of donors and cells, spanning ages from 1 month to 96 years. The data was split into training/validation (train/val) and testing datasets using random donor selection. The train/val dataset comprised 1994 donors with 14,726,170 cells, whereas the testing dataset consisted of 352 donors with 1,504,968 cells (Table S1, Figure 1B-C). The dataset comprised a diverse array of cell types (n=426) sourced from 44 human tissues, including neurons (2,565,212), oligodendrocyte cells (791,513), natural killer cells (333,238), fibroblast cells (161,160), B cells (175,675), and stromal cells (132,596) (Figure 1D, Table S2). Notably, the brain tissue contributed the largest number of cells (5,858,921), followed by other tissues including blood, eye, breast, lung, respiratory system, adipose tissue, kidney, endocrine gland, and skin of body (Figure 1E, Table S3).

### Model training, hyperparameter optimization, and benchmarking

In addition to the gated multi-head attention (GMA) model, scAgeClock also includes multiple alternative models for comparative analysis, including a multi-layer perceptron neural network (MLP), an elastic net regularized linear regression model (Elastic Net), as well as XGBoost-and CatBoost-based models. To assess their performance, each model underwent five-fold cross-validation, with grid-based hyperparameter searching employed to optimize the hyperparameters. For hyperparameter optimization, a random subset of 200,000 age-balanced cells were selected from the original train/val data, while a separate set of 1 million age-balanced cells, also randomly chosen from the train/val data, was used for benchmark model training based on the optimal hyperparameter configuration.

As shown in Figure 2A, which represents one of the training processes for the scAgeClock GMA model, the loss values computed on both the training and validation datasets demonstrated a consistent decreasing trend across the initial training steps. This indicates that the GMA model progressively improved its fit to the data during training. The rate of decrease was more pronounced in the early stages of training and gradually diminished as training progressed. By approximately 1500 training steps, the loss curves for both datasets had largely converged, suggesting that the model had reached a near-optimal state. To evaluate the performance of the scAgeClock GMA model, I first assessed its training process using a grid-searching benchmark dataset. For each hyperparameter combination, the donor-level mean absolute error (MAE) was calculated for each fold of the five-fold cross-validation, and the combination yielding the lowest average MAE across folds was selected as the optimal hyperparameters for the GMA model. The model was then re-trained on the 1 million benchmark train/val dataset using these optimal hyperparameters and evaluated on the independent testing dataset. Correlation analyses revealed a high correlation between chronological ages and predicted ages at both the cell level (Pearson’s r=0.78, Figure 2B) and donor level (Pearson’s r=0.8, Figure 2C).

**Figure 2.**
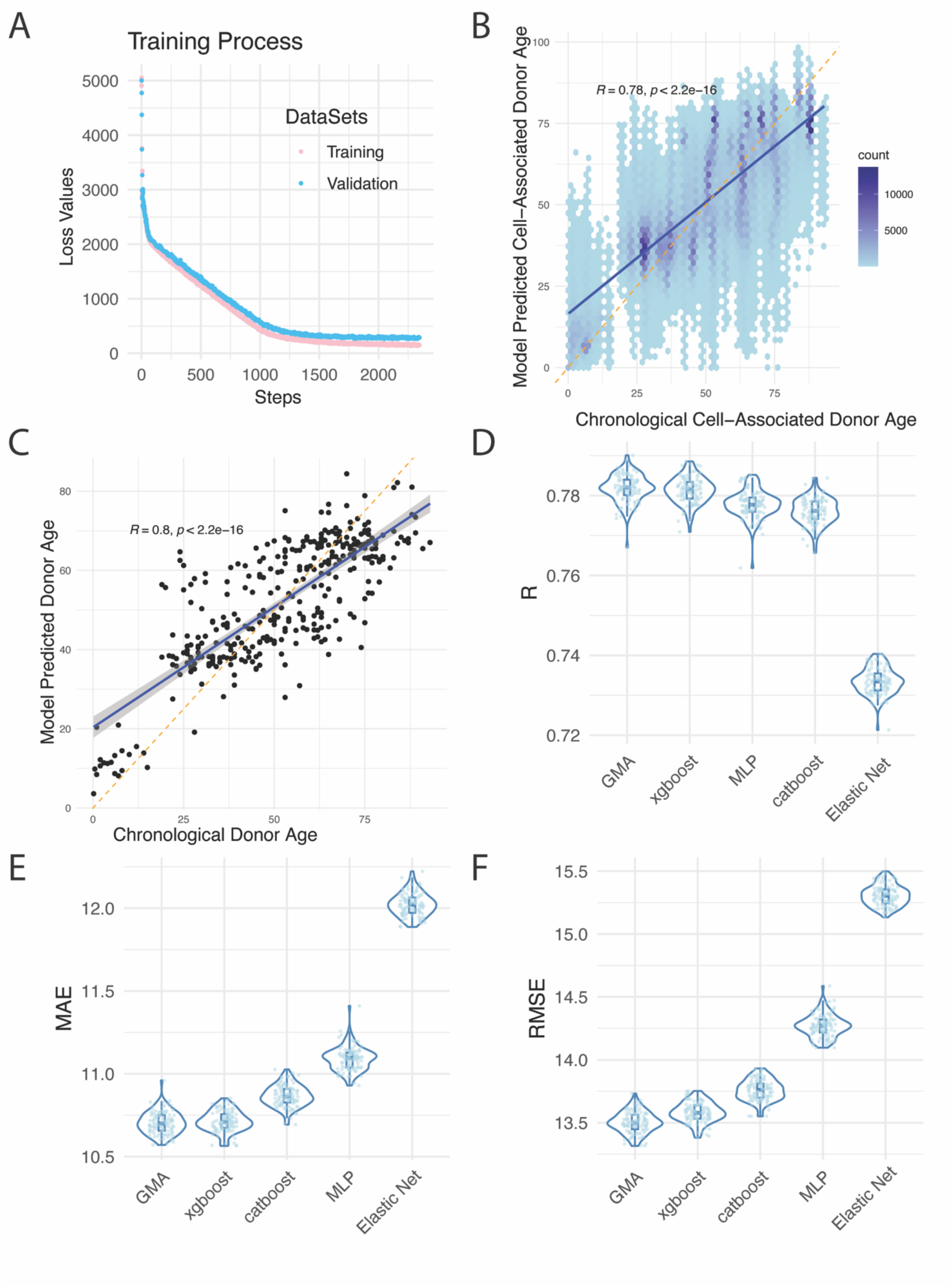
Performance evaluation of scAgeClock GMA model against baseline models. (A) Training and validation loss trajectories for a representative training experiment for scAgeClock GMA model. (B-C) Cell-level (B) or donor-level (C) correlation between chronological age and predicted age by scAgeClock GMA model in the testing dataset. Pearson’s correlation coefficient (R) and associated P-value are shown. (D-F) Comparative performance evaluation of the scAgeClock GMA model against four baseline models - Linear (Elastic Net), Multi-Layer Perceptron (MLP), XGBoost, and CatBoost - using three key metrics: Pearson correlation coefficient (R, D), Mean Absolute Error (MAE, E), and Root Mean Squared Error (RMSE, F).

Next, the optimal hyperparameters for each scAgeClock model were determined through grid searching using the benchmark training and validation dataset. Subsequently, the training and testing procedures were repeated for each model, mirroring the protocol employed for the GMA model. A comparison of model performance on the independent testing dataset revealed that the GMA model achieved the highest Pearson correlation coefficient (Figure 2D) between predicted ages and chronological ages. Specifically, one-sided t-tests indicates that the GMA model outperformed XGBoost, MLP, CatBoost, and Elastic Net with P-values of 0.0026, 3.41e-38, 2.84e-55, and 7.32e-136, respectively. Furthermore, the GMA model exhibited the lowest Mean Absolute Error (MAE, Figure 2E), with one-sided t-tests yielding P-values of 0.017, 1.93e-59, 4.50e-85, and 3.31e-135 against XGBoost, CatBoost, MLP and Elastic Net, respectively. Finally, the GMA model also displayed the lowest Root Mean Squared Error (RMSE, Figure 2F), with one-sided t-tests producing P-values of 5.81e-27, 1.24e-70, 1.32e-106, and 4.48e-144 against XGBoost, CatBoost, MLP, and Elastic Net, respectively.

### Model performance on independent testing dataset across cell types

The scAgeClock GMA model was further trained on the entire train/val dataset using cell-type balanced data loader, initializing from the pre-trained model obtained during benchmarking. Following completion of cell-type level balanced training, the performance of the scAgeClock model was evaluated on the independent testing dataset.

MAE values were calculated for each tissue-level cell type, considering only those with more than 20 cells. A total of 655 tissue-level cell types were evaluated (Figure 3A, Supplementary Table S4), revealing a high degree of diversity in MAE values among these cell types. Notably, 76 tissue-level cell types (11.6%) had MAE values less than 5, while 310 (47.3%), 498 (76.0%), and 604 (92.2%) had MAE values less than 10, 15, and 20, respectively. Eight tissue-level cell types exhibited MAE values less than 3, predominantly from blood and colon tissues (Figure 3B). Conversely, certain tissue-level cell types displayed high MAE values, with cell types from pancreas, liver, and skin tissues among the highest (Figure 3C).

**Figure 3.**
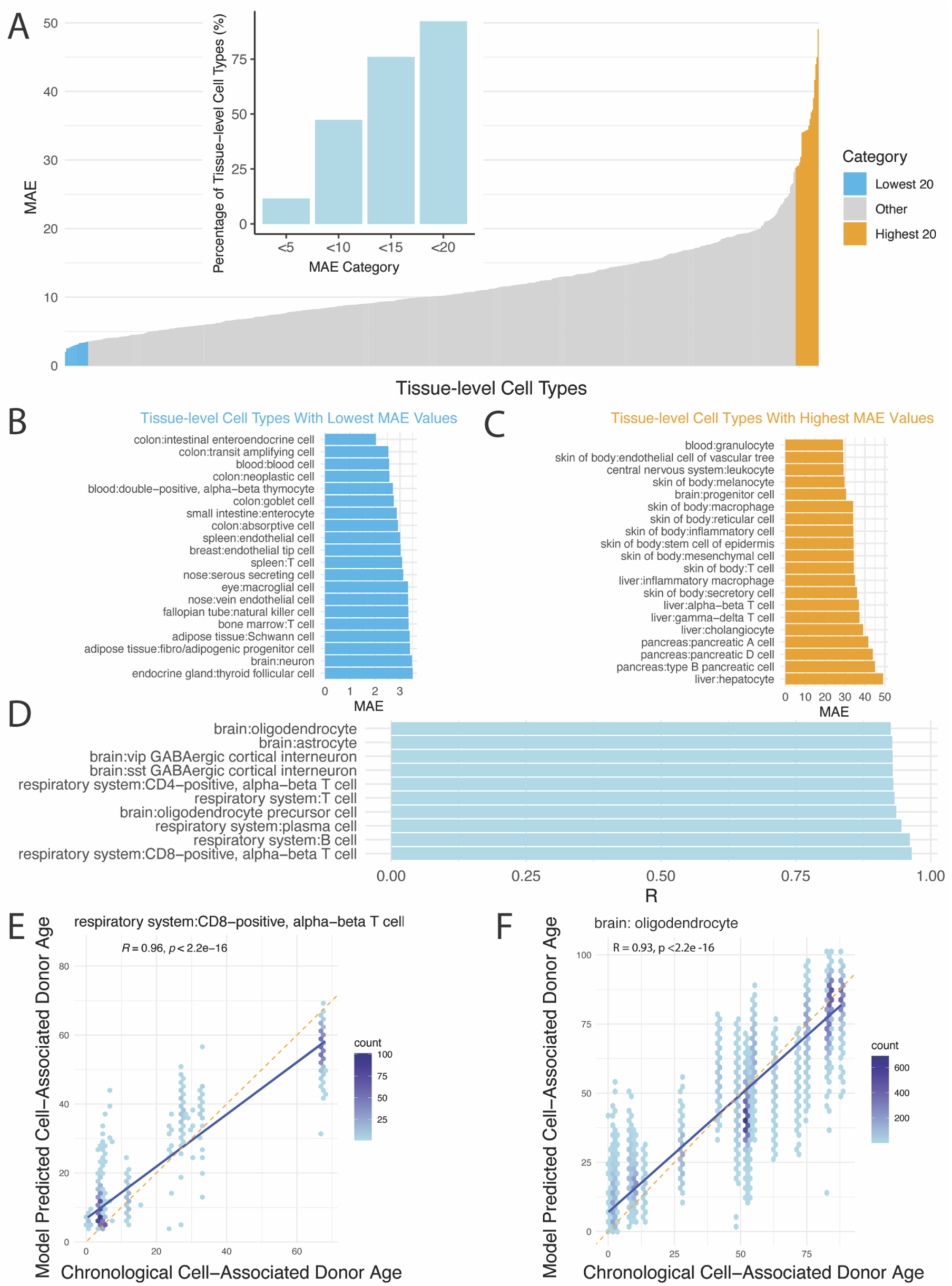
Evaluation of scAgeClock GMA model performance across tissue-level cell types using independent testing datasets. (A) Distribution of MAEs across tissue-level cell types. The lowest 20 and highest 20 cell types were highlighted. (B) Top 20 tissue-level cell types with lowest MAEs. (C) Top 20 tissue-level cell types with highest MAEs. (D) Top 10 tissue-level cell types with highest Pearson correlation coefficients (R). (E-F) Scatter plots of model predicted and chronological ages for two cell types with top correlation coefficients: (E) respiratory system CD8-positive, alpha-beta T cells and (F) brain oligodendrocytes.

Pearson correlation coefficients between model-predicted ages and chronological ages were computed for each tissue-level cell type, considering only those with at least five donors. Seventeen tissue-level cell types exhibited Pearson correlation coefficients (R) greater than 0.9, including respiratory system CD8-positive, alpha-beta T cell (R=0.965), respiratory system B cell (R=0.961), respiratory system plasma cell (R=0.946), brain oligodendrocyte precursor cell (R=0.936), respiratory system T cell (R=0.933), respiratory system CD4-positive, alpha-beta T cell (R=0.931), brain sst GABAergic cortical interneuron (R=0.930), brain vip GABAergic cortical interneuron (R=0.929), brain astrocyte (R=0.929), and brain oligodendrocyte (R=0.926) (Supplementary Table S4 and Figure 3D-F).

### Feature importance and biological interpretation

A total of 19,238 features were utilized for training and evaluating the scAgeClock model, consisting of four categorical features (tissue, cell type, assay and sex) and 19,234 protein-coding genes. To gain insights into the model’s behavior, feature importance was calculated for each feature. Upon ranking by feature importance, it was observed that all categorical features were within the top 10 most important features (Figure 4A and Supplementary Table S5). The top ten genes based on feature importance were identified as RPS17, DEFA5, EEF1G, RPL41, B2M, RPS23, CNTNAP2, RPL26, GPX1, and RPS2 (Figure 4A).

**Figure 4.**
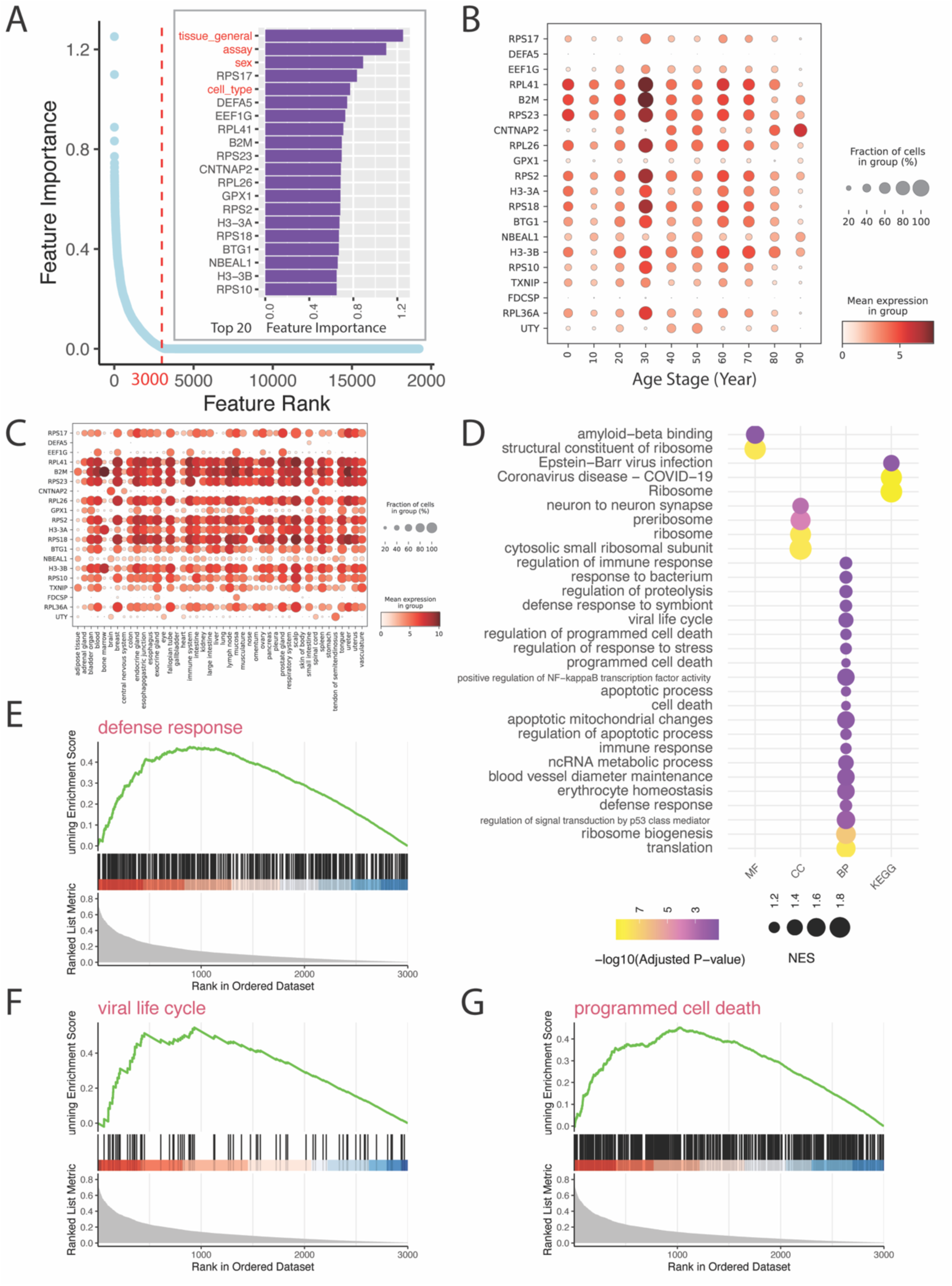
Characterizing feature importance in the scAgeClock GMA model. (A) Feature importance ranking. The top 20 features are shown, with categorical features highlighted. The turning point of feature importance is illustrated by a red dashed line. (B-C) Analysis of gene expression patterns for the top 20 genes ranked by feature importance across different age groups (B) and tissues (C), based on a random sampling of 100,000 age-balanced single-cell transcriptome profiles from the train/val dataset. (D) Gene set enrichment analysis (GSEA) results showing the significantly enriched pathways or Gene Ontology (GO) terms. GSEA was based on the genes within the turning point ranked by feature importance. (E-G) GSEA plots highlighting GO terms related to (E) “defense response”, (F) “viral life cycle”, and (G) “programmed cell death”.

Next, the expression patterns of the top 20 genes with highest feature importances were examined across age stages and tissues. Most of these genes exhibited widespread detection in different age stages and displayed non-linear changes along with ages. Interestingly, out of the top 20 genes, 15 showed peak expression around 30 years old (Figure 4B). Moreover, most of the top genes were also widely expressed across different tissues (Figure 4C).

To investigate the functional enrichment of genes based on feature importance, Gene Set Enrichment Analysis (GSEA) was performed using both Gene Ontology (GO) and Kyoto Encyclopedia of Genes and Genomes (KEGG) databases. Based on the feature importance distribution (Figure 4A), a turning point in the scores was identified around 3000. The GSEA analysis was performed using the top 3000 genes ranked by their feature importance scores. Notably, the ribosome-related genes were highly enriched among the top feature importance genes, with significant enrichment P-values in both KEGG Ribosome pathway (Benjamini-Hochberg or BH adjusted P-value=2.1e-9) and GO ribosome terms (BH adjusted P-value=1.5e-8) (Figure 4D, Supplementary Table S6). This finding is consistent with a previous aging clock study based on the blood samples of supercentenarians, which highlighted the importance of ribosome-related genes during the aging process and proposed that those genes might be related to low inflammation and slow aging [11]. Furthermore, genes related to “defense response” (Figure 4E), “viral life cycle” (Figure 4F), and “programmed cell death” (Figure 4G) were also found to be enriched with top feature importance genes, with BH-adjusted P-values of 0.01, 0.04, and 0.03, respectively.

### Aging deviation analysis of human cell types

To investigate the aging deviation between model-predicted ages and chronological ages for various human cell types, this study introduces the Aging Deviation Index (ADI), which was calculated for each tissue-level cell type. The ADI ranges from 0 to 1, with higher scores indicating greater age acceleration and lower scores suggesting greater age deceleration. Notably, significant heterogeneity was observed in the ADI distribution across different tissue-level cell types (Figure 5A). Specifically, liver hepatocytes, pancreas pancreatic cells, liver gamma-delta T cells, and brain progenitor cells exhibited the lowest ADIs (Figure 5B), whereas cell types from skin tissues displayed the highest ADIs (Figure 5C).

**Figure 5.**
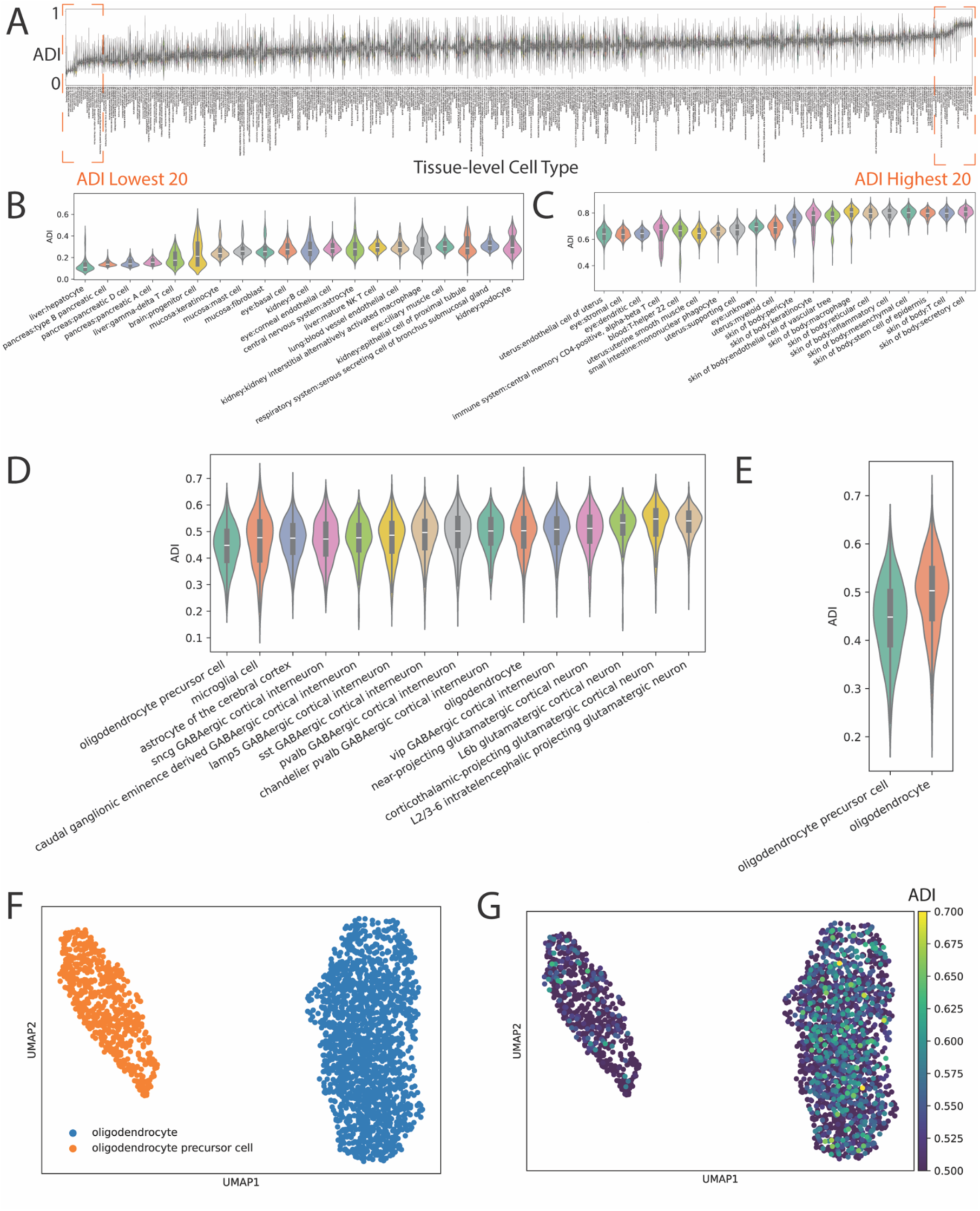
Aging Deviation Index (ADI) analysis of human cell types. (A) Overview of the ADI distribution across human cell types (tissue-level). (B-C) The top 20 tissue-level cell types with (B) lowest and (C) highest ADIs. (D-G) Intra-donor comparison of ADIs among cell types. The donor (donor id: H21.33.019) with the largest number of brain cells in the testing dataset was selected. (D) ADI comparisons across brain cell types, (E) Comparison of ADIs between oligodendrocyte precursor cells (OPCs) and oligodendrocytes (OLs), (F) UMAP visualization of OPCs and OLs, and (G) UMAP visualization of ADIs among OPCs and OLs.

To further examine the differences in ADI among cell types within an individual, this study focused on the donor in the testing dataset with the largest number of brain cells. As shown in Figure 5D, on average, oligodendrocyte precursor cells had the lowest ADIs compared to other cell types, while glutamatergic neurons exhibited the highest ADIs.

To explore the relationship between ADI and cell differentiation potency, two cell types with established differentiation relationships in the human brain were analyzed: oligodendrocytes (OLs) and oligodendrocyte precursor cells (OPCs). As depicted in Figure 5E-G, OPCs had significantly lower ADIs compared to OLs on average, with a P-value of 1.03e-40 based on the one-sided Wilcoxon rank-sum test.

### Transcriptomic aging acceleration in human diseases revealed by scAgeClock

To investigate the disease-related changes in transcriptomic biological age, the trained scAgeClock GMA model was applied to an independent human cortex dataset from the ZEBRA Brain Atlas[22]. This comprehensive repository contains human brain scRNA-seq datasets from various diseases, including Autism Spectrum Disorder (ASD), COVID-19, Multiple Sclerosis (MS), Frontotemporal Dementia (FTD), Alzheimer’s Disease (AD), Huntington’s Disease (HD), and normal control (CT),.

To examine age acceleration in disease-associated cells, the scAgeClock ADI score was calculated for each cell. The ADIs were compared between age-matched healthy controls and diseased groups. Among 18 comparisons, 16 showed significantly lower average ADIs in cells from healthy control groups than in those from diseased groups (Figure 6A), with one-sided Wilcoxon test P-value less than 0.01. These finding suggest that the aging clock of disease-associated cells may be aberrantly accelerated compared to age-matched normal cells.

**Figure 6.**
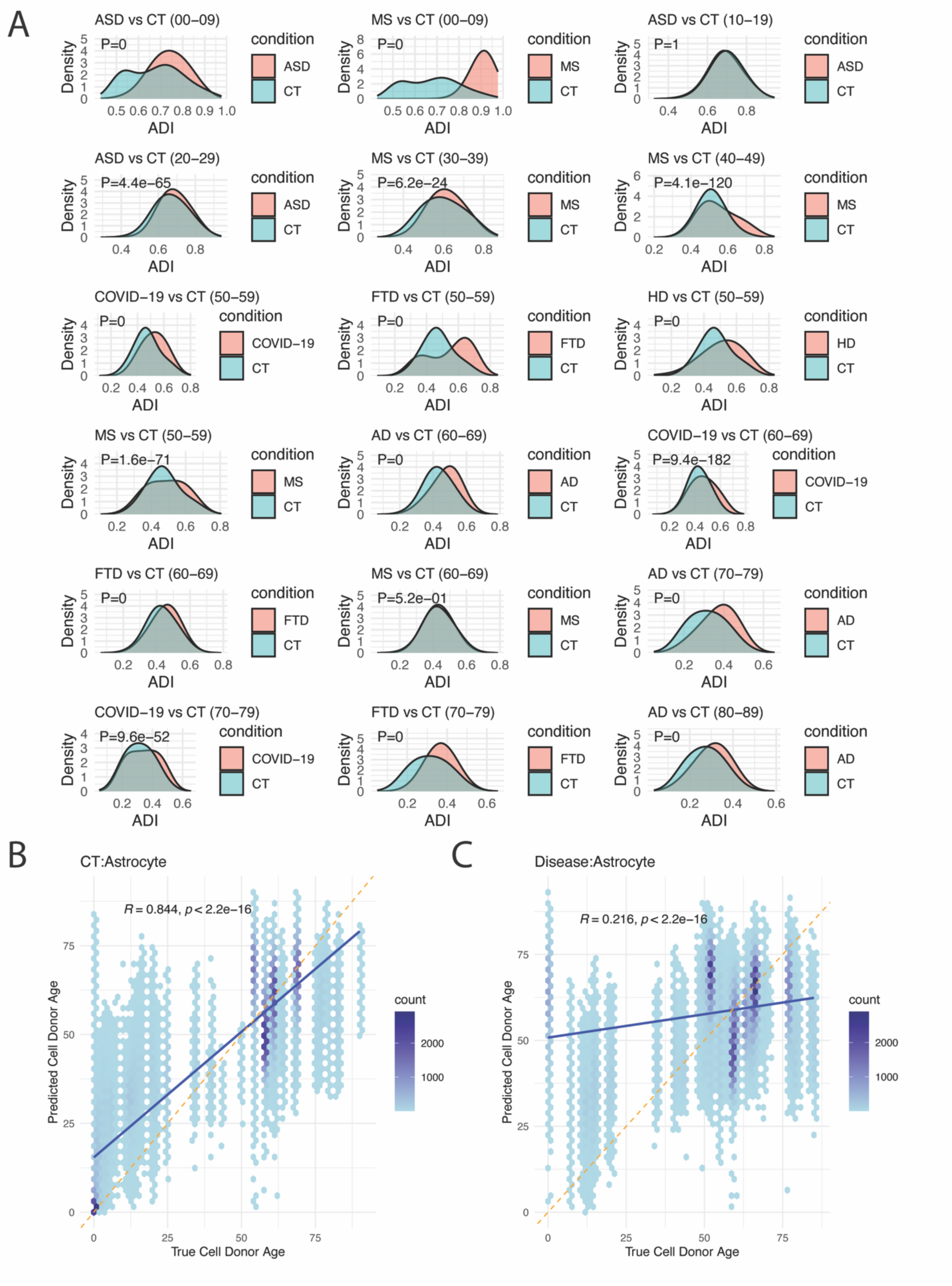
Comparative analysis of ADIs in human diseases and healthy controls. (A) Single cell level ADI distribution comparison between disease and normal conditions. CT, normal control; FTD: Frontotemporal Dementiam; AD: Alzheimer’s Disease; ASD, autism spectrum disorder; MS, multiple sclerosis; and HD, Huntington’s disease. Donor age ranges are provided in parentheses. P-values from one-sided Wilcoxon tests (CT group < disease group) are displayed in each sub-plot. (B-C) Scatterplots illustrating the relationship between chronological ages and predicted ages for human brain astrocytes in normal control (B) or disease group (C). The disease group encompasses all the disease conditions listed in (A). Pearson’s correlation coefficient (R) and associated P-value are shown for each plot.

Higher Pearson’s correlation coefficients between scAgeClock-predicted ages and chronological ages were observed in cells from healthy groups than those from diseased groups (Figure 6B-C, Supplementary Figure S1). Notably, astrocytes from healthy controls exhibited a high Pearson’s correlation coefficients of 0.844 between predicted ages and chronological ages (Figure 6B), whereas this value was significantly lower (0.216) for the same cell types from diseased groups (Figure 6C). Similarly, higher correlations between scAgeClock-predicted ages and chronological ages were also observed in neurons (Supplementary Figure S1A-D) and microglia cells (Supplementary Figure S1E-F) from healthy donors compared to the diseased groups.

### Transcriptomic aging acceleration in higher tumor phases and metastatic tumor cells

Cancer cells are characterized by uncontrolled growth and division, as opposed to aging cells, which lose proliferative capacity over time. To investigate the potential application of scAgeClock’s ADI in cancer research, a human pan-cancer scRNA-seq dataset was obtained from a recent study [23]. This analysis revealed that cells from seven distinct cancer types exhibited lower ADIs in higher tumor phases (Figure 7A). Specifically, this trend was observed in acral lentiginous melanoma (ALM), combined hepatocellular-cholangiocarcinoma (CHC), kidney renal clear cell carcinoma (KIRC), skin cutaneous melanoma (SKCM), head and neck squamous cell carcinoma (HNSC), nasopharyngeal carcinoma (NPC), and pancreatic adenocarcinoma (PAAD). For instance, the average ADI of cells from phase IIIB ALM was significantly higher than that cells from phase IV ALM (one-sided Wilcoxon test P-value=0). Similarly, the average ADI of cells from phase IB PAAD was also significantly higher than that of cells from phase IVB PAAAD (one-sided Wilcoxon test P-value=0).

**Figure 7.**
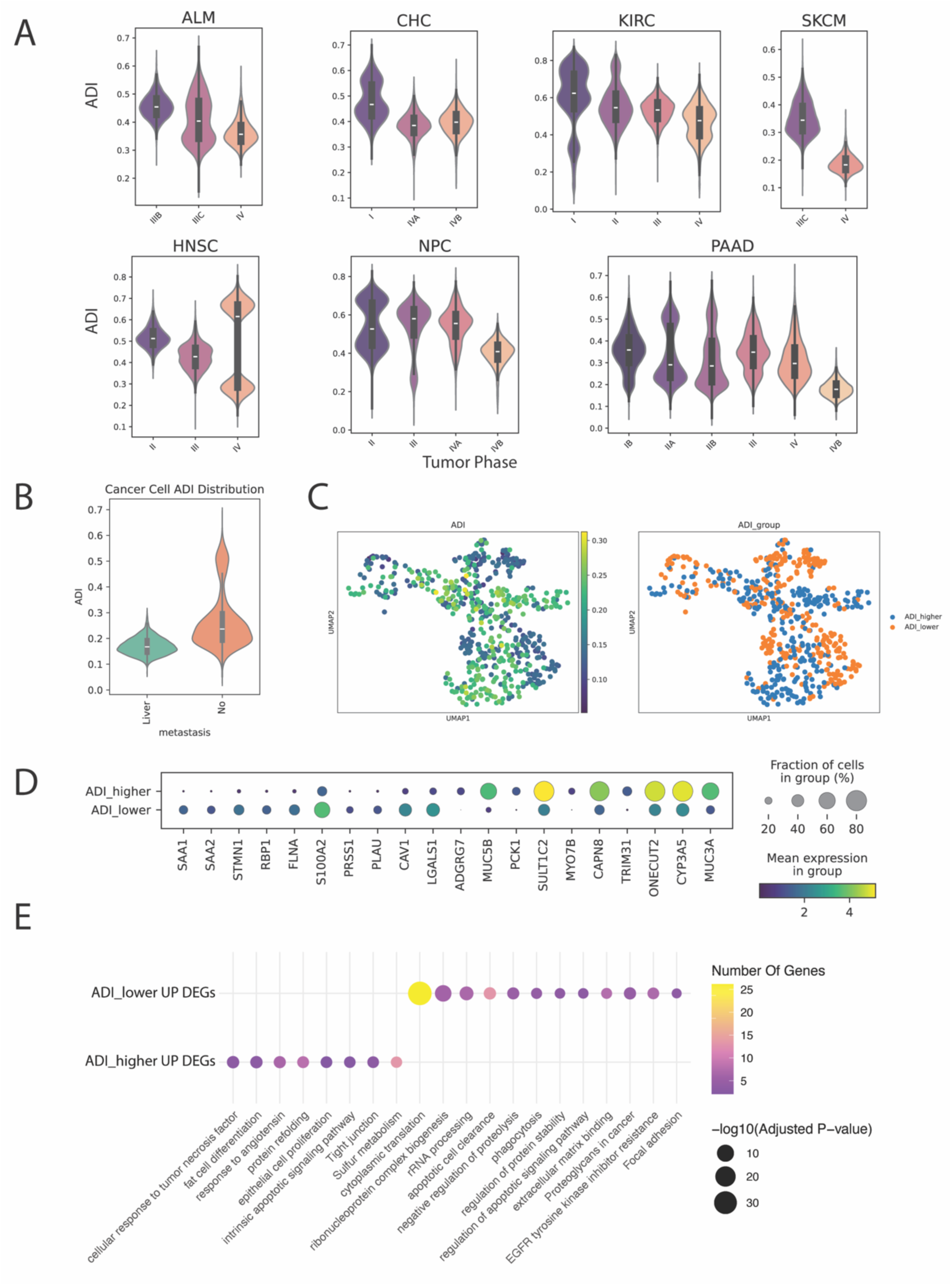
Tumor heterogeneity and metastasis analysis using ADI. (A) ADI distribution across tumor phases in various human cancer types. Comparative analysis of ADI distribution among different tumor phases in acral lentiginous melanoma (ALM), combined hepatocellular-cholangiocarcinoma (CHC), kidney renal clear cell carcinoma (KIRC), skin cutaneous melanoma (SKCM), head and neck squamous cell carcinoma (HNSC), nasopharyngeal carcinoma (NPC), and pancreatic adenocarcinoma (PAAD). (B) ADI comparison between metastatic and primary tumor cells in PAAD. ADI distribution analysis of PAAD tumor cells from metastatic liver tissue versus primary tumor site. (C) UMAP visualization of ADI-stratified PAAD primary tumor cells. Uniform Manifold Approximation and Projection (UMAP) representation of PAAD primary tumor cells stratified into ADI-higher and ADI-Lower groups. (D) Differentially expressed genes (DEGs) between ADI-higher and ADI-lower PAAD primary tumor cells. (E) Enriched KEGG pathways and GO terms in ADI-stratified PAAD primary tumor cells.

To further dissect the characteristic of tumor cells, a PAAD donor with the largest number of cells was selected for in-depth ADI analysis. Notably, ADIs of tumor cells from metastatic PAAD tissues were significantly lower than those from primary PAAD tissues (Figure 7B, one-sided Wilcoxon test P-value=2.3e-23). To explore the functional implications of ADI heterogeneity within primary tumor cells, two subgroups of tumor cells with higher ADIs (one standard deviation above the mean, ADI-higher) and lower ADIs (one standard deviation below the mean, ADI-lower) were identified for further investigation (Figure 7C).

Differential gene expression analysis between ADI-higher and ADI-lower PAAD primary tumor cells revealed a total of 719 differentially expressed genes (DEGs), with 153 genes up-regulated in ADI-lower tumor cells and 566 genes up-regulated in ADI-higher tumor cells (Supplementary Table S7). The top up-regulated DEGs in the ADI-lower group included serum amyloid A1 (SAA1), serum amyloid A2 (SAA2), stathmin 1 (STMN1), retinol binding protein 1 (RBP1), and filamin A (FLNA). In contrast, the top up-regulated DEGs in the ADI-higher group included adhesion G protein-coupled receptor G7 (ADGRG7), mucin 5B, oligomeric mucus/gel-forming (MUC5B), phosphoenolpyruvate carboxykinase 1(PCK1), sulfotransferase family 1C member 2(SULT1C2), and myosin VIIB(MYO7B) (Figure 7D).

Functional enrichment analysis of the DEGs revealed that ADI-lower PAAD primary tumor cells were enriched with genes involved in “rRNA processing”, “phagocytosis”, “Proteoglycans in cancer”, “focal adhesion”, and “regulation of protein stability”. In contrast, ADI-higher PAAD primary tumor cells were enriched with genes involved in “fate cell differentiation”, “cellular response to tumor necrosis factor”, “protein refolding”, “response to angiotensin”, and “intrinsic apoptotic signaling pathway” (Figure 7E).

## DISCUSSION

There are two distinct types of ages: chronological age, which measures the time elapsed since birth, and biological age, which reflects the physiological status of our body. Aging clock research on supercentenarians has shown that these individuals tend to have younger biological ages compared to their chronological ages[11]. Genetic studies have also identified variants associated with human longevity (reviewed by [24, 25]), including those in the apolipoprotein E (APOE) gene[26, 27]. Notably, APOE is among the top 3% of genes ranked by the feature importance scores from scAgeClock (420 out of 19234 protein coding genes used in the model), further supporting its role in the aging process. In contrast, studies on patients with accelerated aging diseases, such as Hutchinson-Gilford progeria syndrome, have implicated genetic changes in human lamin A/C (LMNA) [28, 29]. Interestingly, LMNA is also among the top 3% of most important genes for scAgeClock model (436 out of 19234), with a feature importance like that of APOE. However, more than 400 genes that have higher rankings than LMNA and APOE based on scAgeClock’s feature importances (Supplementary Table S5), suggesting a complex interplay of gene networks involved in the human aging process.

Indeed, gene set enrichment analysis revealed that 97 GO terms are significantly (BH-adjusted P value less than 0.05) enriched with genes ranking highly by scAgeClock, including those related to programmed cell death(Adjusted P value = 0.029), ribosome (Adjusted P value = 1.52e-08), translation (Adjusted P value = 1.52e-08), positive regulation of cell growth (Adjusted P value = 0.0075), erythrocyte homeostasis (Adjusted P value = 0.011), apoptotic process (Adjusted P value = 0.025), regulation of response to stress (Adjusted P value = 0.035), immune response (Adjusted P value = 0.021), and blood vessel diameter maintenance (Adjusted P value = 0.019) (Table S6). Collectively, these results underscore the significant role that these biological processes play in modulating the aging process.

Aging clock models have been employed not only to identify genes associated with aging but also as biomarkers to assess the efficacy of anti-aging interventions. Currently, approximately 15 registered clinical trials for anti-aging interventions have been initiated or completed, with most utilizing DNA methylation-based aging clock models as outcome measurement[30]. These models primarily rely on DNA methylation data from blood samples and have reported MAE values ranging from 3 to 5 years[3]. In contrast, current published transcriptome-data based aging clock models, also mainly derived from blood samples, exhibit higher age prediction errors compared to their DNA methylation-based counterparts (MAEs of 5-7 vs 3-5 years)[3]. This disparity in prediction accuracy may be attributed to the cell heterogeneity of gene expressions, which has a lesser impact on DNA methylation-based models. This study reveals that the scAgeClock model performs with varying degrees of accuracy across different cell types, yet the best-performing cell types achieve lower MAE values (2.03 – 3.50 years; Figure 3B) than previous DNA methylation-based aging clock models. Notably, it has been suggested that an aging clock model capable of perfectly predicting chronological age may not necessarily be optimal for biological age prediction and its applications[3]. The cell type-specific variations in age prediction observed with scAgeClock may indicate differences in aging rates at the cellular level, which can be further investigated through experimental studies in the future.

This study also introduces the concept of an Aging Deviation Index (ADI), a quantitative measure that ranges from 0 to 1. The ADI provides a novel way to assess the likelihood of accelerated or decelerated aging, with higher values indicating a greater probability of accelerated aging and lower values suggesting a slower pace of aging. Notably, this study has demonstrated that ADIs can serve as a proxy measurement for various cellular processes, including cell disease status, cell differentiation potential, and cancer cell malignancy degree. The ADI offers a promising avenue for exploring new applications based on scAgeClock model. Future studies may investigate the use of ADIs in evaluating the efficacy of anti-aging interventions, enabling early disease predictions, and facilitating patient stratification in clinical trials. By leveraging the ADI, researchers and clinicians may gain valuable insights into the aging process and develop more effective strategies for promoting healthy aging and preventing age-related diseases.

Although the scAgeClock model was trained on an extensive dataset comprising over 10 million single-cell transcriptome profiles, there is still room for improvement. Notably, some cell types or tissues are under-represented in the training data (Figure 1D). For instance, cells from brain tissues are over-represented, while certain human tissues such as testis are not covered by the current dataset (Figure 1E). Furthermore, there is an age-related imbalance in the distribution of cells, with fewer cells from teenage stages and longevity groups (> 100 years) represented in the training data (Figure 1C). To address these imbalances, scAgeClock features a balanced data loader that enables balanced sampling of cells from each label or feature. This ensures that the model is trained on a representative dataset, mitigating potential biases. Moreover, scAgeClock allows for versatile piped training, where the model can be initially trained using an age-balanced data loader and then fine-tuned using a cell-type-balanced data loader. The continued advancement of single-cell RNA sequencing technologies is expected to yield a vast increase in human scRNA-seq datasets in the coming years. This will provide opportunities for further improving the aging-clock prediction performance of scAgeclock, enabling more accurate and reliable predictions across diverse cell types, tissues, and age groups.

In conclusion, this study presents an important contribution to the field of aging research with the development of a novel gated multi-head attention-based neural network architecture, GMA, and its implementation in the open-sourced scAgeClock software. By leveraging over 10 million single-cell transcriptome profiles from 44 human tissues and over 400 human cell types, scAgeClock provides an unparalleled platform for aging clock modeling. Furthermore, the integration of multiple benchmark methods, including Elastic Net, XGBoost, CatBoost, and MLP, facilitates seamless comparisons and evaluations by the aging research community. Notably, this study also introduces the concept of an Aging Deviation Index (ADI), which is generated by scAgeClock and has demonstrated its potential applications in various fields. The ADI offers a novel metric for assessing the likelihood of accelerated or decelerated aging and its implications on cellular process, disease progression and treatment outcomes. Overall, scAgeClock represents a valuable tool for the aging research community, providing a comprehensive platform for investigating the complex mechanisms underlying aging and age-related diseases. By making this software openly available, it is expected to accelerate the pace of discovery in this filed and ultimately contribute to the development of effective interventions for promoting healthy aging.

## METHODS

### Data Collection

The CZ CELLxGENE human single-cell RNA sequencing (scRNA-seq) datasets, including raw sequencing counts and associated cell-level metadata annotations, were retrieved using the CELLxGENE Census Python API (https://chanzuckerberg.github.io/cellxgene-census/python-api.html). Specifically, the cellxgene-census Python package was employed to access the API, and only primary data records without duplicates were considered by setting ‘is_primary_data=Truè. The ‘get_anndatà function from cellxgene-census tool was utilized for downloading the datasets, with the organism parameter set to “Homo sapiens”. The downloaded data were then stored in .h5ad format files. To facilitate large-scale downloading, the entire metadata table was first retrieved, followed by splitting the data records based on the assay column. Subsequently, each assay’s data was downloaded independently. Verification of the completeness of the data downloaded from CZ CELLxGENE was performed by cross-referencing the downloaded records with the metadata table using the cell IDs (soma_joinid column).

To investigate the model’s performance in tissue-level cell types, the whole CZ CELLxGENE testing dataset was used. To further explore the model’s performance on brain oligodendrocyte precursor cells (OPCs) and oligodendrocytes (OLs), the normal donor (donor id: H21.33.019) with the largest number of brain cells from testing dataset was selected.

To evaluate the performance of the scAgeClock model in the context of various human brain diseases, raw count datasets along with associated metadata annotations for the human ZEBRA brain atlas [22] were obtained from the ZEBRA website (https://ccb-compute.cs.uni-saarland.de/brain_atlas/). The ZEBRA database comprises scRNA-seq datasets from both human cortex and non-cortex tissues. Specifically, the cortex tissues include “Temporal cortex”, “Occipital cortex”, “Frontal cortex”, “Prefrontal cortex” “Cortex”, “Cingulate cortex”, and “Parietal cortex”. In contrast, the non-cortex tissues encompass “Substantia nigra”, “Caudate nucleus”, “Hippocampus”, “Choroid”, “Putamen”, “Nucleus accumbens”, “Cerebellum”, “Spinal cord”, and “Olfactory bulb”. Due to the greater heterogeneity of non-cortex tissues compared to cortex tissues, focus was placed on the ZEBRA human cortex tissues for cross-conditions comparison (diseases vs normal).

To examine the model’s application on cancer, the human pan-cancer scRNA-seq dataset was downloaded from Zenodo (https://zenodo.org/records/15554080), which was based on a recent human cross-tissue cancer-related single-cell transcriptomic study [23].

### Filtering and splitting of scRNA-seq data

The primary human CZ CELLxGENE data was further filtered through a multi-step process for model construction and evaluation. Firstly, the data was filtered by disease type (disease column in the metadata), retaining only normal samples. Next, the data was selected based on accurate age information (development_stage column in the metadata), with all ages converted to years. The data was then filtered by sequencing technology (assay column), considering only the assays with more than 1,000 cells. Subsequently, the data was filtered by cell type, selecting only those with more than 200 cells. Finally, the data was filtered by donor (donor_id column), considering only donors with more than 1,000 cells. Notably, prior to filtering by donor, sex and age information were appended to the original donor IDs for certain donors to account for duplicate IDs.

To enable further analysis, the processed human normal CZ CELLxGENE datasets were split into two independent parts: one for training and validation (train/val) and another for independent testing. The number of cells for each donor was calculated, and the top four donors with the highest cell counts were assigned to the train/val dataset. From the remaining donors, 15% were randomly selected as testing datasets, while the rest were included in the train/val dataset. Furthermore, to facilitate cross-validation, the donors within train/val datasets were further randomly divided into five equal folds.

### Feature selection and data preprocessing

Following cell filtering, the human normal CZ CELLxGENE data underwent gene expression quantification. For each gene, the number of cells with detectable expression was calculated, where a gene was considered expressed if it had at least 1 unique molecular identifier (UMI) counts. To select relevant genes for analysis, only those expressed in at least 1,000 cells were retained. To further reduce the dimensionality of the input data, only protein-coding genes were selected for modeling. The identification of protein-coding genes was based on the “Gene type” annotation from the Ensembl Biomart database (Genome version GRCh38.p14). Next, gene expression levels based on UMI counts in each cell underwent normalization and scaling using the scanpy.pp.normalize_total (target_sum=1e6, Couts Per Million Normalization or CPM) and scanpy.pp.log1p (default settings) functions from scanpy[31].

### Data formatting and dataloader construction

Four categorical features from CZ CELLxGENE were selected for modeling: assay, tissue (tissue_general column in the metadata), sex, and cell type (cell_type column in the metadata). For each categorical feature type, string values were sorted in ascending order and then assigned a numeric value based on their index after sorting, starting from 0. The resulting numeric values for the categorical features were then concatenated with the gene expression matrix as the first four columns, in the order of assay, cell type, tissue, and sex.

To facilitate modeling, scAgeClock provides a dataloader class (H5ADDataLoader) for iterating over batch-level data in .h5ad format input files. To reduce memory usage, scAgeClock leverages scanpy’s backed mode loading feature, which stores the entire dataset in disk file rather than loading it into memory.

Additionally, scAgeClock provides a balanced dataloader (BalancedH5ADDataLoader) for retrieving balanced batches of data based on a given type of cell attributes such as cell type, tissue, and age category. Given a batch size N and M unique cell attributes, a mini-batch size 𝑁_𝑚𝑖𝑛𝑖_ is defined as 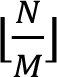. During each batching iteration, BalancedH5ADDataLoader retrieves 𝑁_𝑚𝑖𝑛𝑖_ data from attribute type. The remaining 𝑁_𝑟𝑒𝑚𝑎𝑖𝑛𝑑𝑒𝑟_ = 𝑁 𝑚𝑜𝑑 𝑀 data are randomly sampled from the unsampled cells.

### Gated multi-head attention (GMA) neural networks

The scAgeClock GMA neural networks comprise five primary components: (1) a categorical feature embedding layer; (2) a regularized feature gating layer; (3) a dimensional reduction feature projection layer; (4) a multi-head attention layer; and (5) a fully connected neural network layer (FC) with age prediction output.

The scAgeClock neural network architecture is built using PyTorch (https://pytorch.org), where input data and intermediate computations are represented as tensors, multidimensional matrices containing the original or transformed data. Initially, categorical features are embedded using ‘torch.nn.Embedding’ function with an embedding dimensionality of 4, which is then concatenated with the gene expression tensor.

To filter out irrelevant features, the concatenated inputs tensor passes through a feature gating neural network, which employes an elastic net regularized linear transformation. The gating layer includes two hyperparameters, ‘l1_lambdà and ‘l2_lambdà, to control the weights for L1 (Lasso) and L2 (Ridge) regularization, respectively.

Next, to reduce the dimensionality of the gated tensor, a feature projection neural network layer is applied, which performs an affine linear transformation on the incoming tensor with a smaller output dimension size (default value = 512).

Subsequently, the dimension-reduced tensor undergoes further transformation by a multi-head attention neural network based on ‘torch.nn.MultiheadAttention’, with 8 default heads. Finally, the output from the multi-head attention is fed into an age prediction neural network layer, which consists of multiple hidden layers (default hidden layer sizes: [256,128]) and one output layer. The output layer generates the age prediction. Additionally, to introduce nonlinearity between the hidden and output layers of the FC neural network, a ReLU (Rectified Linear Unit) activation function is applied.

### Multi-layer perceptron neural networks (MLP) and the elastic net linear regression (Elastic Net) model

Like the scAgeClock GMA model, both MLP and Elastic Net models are built using PyTorch. The inputs are transformed by an embedding neural network layer, consistent with the GMA model. In the MLP neural networks, the embedded input tensor is then processed through multiple hidden layers, sequentially applying ReLU non-linear transformation and dropout processing. By default, scAgeClock MLP employs three hidden layers with 512 (first), 256 (second), and 128 (third) neurons, respectively, and a dropout probability of 0.2. For the Elastic Net, the embedded input tensor is further transformed by an affine linear regression with elastic net regularized weights added to the loss function. Like the feature gating layer in GMA model, the Elastic Net model comprises two hyperparameters, ‘l1_lambdà and ‘l2_lambdà, which control the weights for L1 (Lasso) and L2 (Ridge) regularization, respectively. Ultimately, both the MLP and Elastic Net models produce age predictions through their final output layers.

### XGBoost and CatBoost Models

The XGBoost and CatBoost models in scAgeClock are built on top of the publicly available Python packages xgboost (https://xgboost.readthedocs.io/en/stable/install.html) and catboost (https://catboost.ai/docs/en/concepts/python-installation), respectively. Both gradient boosting models have been integrated into scAgeClock with customized data loaders and training modules. Specifically, in the scAgeClock CatBoost dataloader, categorical column indices are passed to the CatBoost Pool constructor, enabling CatBoost to treat these categorical features as label-encoded data.

### Optimization of model training hyperparameters and benchmark comparison

To optimize hyperparameters for each model, grid-based searches were conducted using randomly selected and age-balanced benchmark datasets. Given the computational cost (for all the models) and memory requirements (for the gradient boosting models), the using of benchmark data allowed efficient exploration of the hyperparameter space. The age-balanced benchmark datasets of different sizes were generated based on five-fold split train/val datasets. A smaller dataset comprising 200,000 cells was used for hyperparameter grid searching, while a larger dataset consisting of 1 million cells was employed for training the model with the optimal set of hyperparameters. To ensure an age-balanced dataset, an ‘age stage’ column was introduced into the metadata, which categorized donors into 10-year intervals (e.g., age stages 0-9, 10-19, etc.) using floor division of the ‘age’ values. For each of the five folds of train/val data, the H5ADSampler tool from scAgeClock was utilized to randomly sample age-balanced cells. The sampling process was conducted with the following parameter settings: ‘meta_category_colname=’age_stage’, category_balanced=True, sample_replace=Falsè. The number of sampled cells for each fold was dependent on the specific application, with 40,000 cells being sampled for hyperparameter grid searching and 200,000 cells being sampled for training the model under the optimal set of hyperparameters. This tool employs a similar algorithm as the BalancedH5ADDataLoader from scAgeClock to sample balanced data based on age stage labels.

For the PyTorch-based models (GMA, MLP, Elastic Net), a grid search was performed over three hyperparameters: ‘learning_ratè, ‘l1_lambdà, and ‘l2_lambdà. The value ranges for these hyperparameters were ‘[0.01, 0.001, 0.0001]’ for learning rate, ‘[0.1,0.5]’ for L1 regularization strength (l1_lambda), and ‘[0.1,0.5]’ for L2 regularization strength (l2_lambda). For the gradient boosting-based models (XGBoost, CatBoost), a grid search was performed over two hyperparameters: ‘learning_ratè and ‘boost_depth’, with value ranges of ‘[0.1,0.01,0.001]’ and ‘[6,8]’, respectively. For each combination of hyperparameters, the model was trained using five-fold cross-validation. To facilitate this training process, a ‘training_pipelinè function from scAgeClock was created, providing a unified interface for the training of all scAgeClock models. Notably, for PyTorch-based models, the NVIDIA CUDA device (NVIDIA GeForce RTX 4090, CUDA Version: 12.8) was utilized to accelerate the training process, whereas only CPU devices were used for boosting-based models due to high memory requirement.

After completing the grid search-based model training with five-fold cross-validation, the average validation metrics were computed to facilitate hyperparameter selection. Specifically, the Mean Absolute Error (MAE) was calculated for each hyperparameter setting in each cross-validation fold based on the donor-level age prediction results. The donor-level predicted age was obtained by averaging the corresponding cell-level predicted ages. Subsequently, for each hyperparameter setting, the average donor-level MAE values were computed across all folds of the cross-validation process. Finally, the best hyperparameters for each model were identified as those yielding the minimum average donor-level MAE value.

To facilitate the comparison of different scAgeClock models, the best-performing hyperparameters were used to train each model on 1 million age-balanced benchmark cells randomly sampled from the entire train/val dataset. The trained models were then employed to generate age predictions on the independent benchmark testing dataset. Subsequently, three performance metrics – Mean Absolute Error (MAE), Root Mean Squared Error (RMSE), and Pearson’s correlation coefficient (r) – were computed for each model based on its age prediction performance on the independent testing dataset.

### Model training

For the PyTorch-based models (GMA, MLP, Elastic Net), similar training pipelines were employed. The Mean Squared Error (MSE) loss function was used to calculate the neural network’s loss using PyTorch’s ‘torch.nnMSELoss’ function, while Adaptive Moment Estimation (Adam) served as the optimization algorithm via PyTorch’s ‘torch.optim.Adam’ function. The ‘BasicDataLoader’ from scAgeClock was utilized as the data loader for training the neural networks in batches, with a default batch size of 1,024. Additionally, weights-based penalties were incorporated into the loss function for Elastic Net and GMA models. Specifically, the regularization penalty consisted of combined L1 and L2 penalties, defined as ‘𝑙𝑜𝑠𝑠_𝑟𝑒g_ = 𝑙1_𝑙𝑎𝑚𝑏𝑑𝑎 × |𝑤|_1_ + 𝑙2_𝑙𝑎𝑚𝑏𝑑𝑎 × ‖𝑤‖_2_^2^’. Here, ‘w’ represents the weight vector ‘[𝑤_1_, 𝑤_2_, . . ., 𝑤_𝑁_]’, where ‘N’ is the total number of features, including any embedded features. For Elastic Net, weights were based on the magnitudes of all linear coefficients, whereas for GMA, they were derived from the gating layer’s neuron weights. The contribution of L1 and L2 penalties could be adjusted via hyperparameters ‘l1_lambdà and ‘l2_lambdà, respectively. For models with regularization penalties (Elastic Net and GMA), the total loss was defined as ‘𝑙𝑜𝑠𝑠_𝑡𝑜𝑡𝑎𝑙_ = 𝑙𝑜𝑠𝑠_𝑀𝑆𝐸_ + 𝑙𝑜𝑠𝑠_𝑟𝑒g_’. Furthermore, early stopping during each training epoch was implemented by configuring the ‘train_batch_iter_max’ hyperparameter in the PyTorch-based aging clock.

For the boosting-based models (XGBoost and CatBoost), the native training functions from their respective Python packages are used. The default loss functions employed by these packages are also adopted as the default loss functions for the scAgeClock boosting-based models, specifically Mean Squared Error (MSE) for XGBoost and Root Mean Squared Error (RMSE) for CatBoost. Additionally, custom data loaders are created to cater to the specific needs of the boosting-based models within scAgeClock, namely ‘CatBoostDataLoader’ and ‘XGBoostDataLoader’. These specialized data loaders enable scAgeClock to provide a batch-level training interface, streamlining the model training process.

For the GMA model, scAgeClock offers ‘initial_model’ option to enable training from a pre-trained model. In addition to benchmarking the GMA model on benchmark datasets for comparison, it was also trained on the full train/val datasets. The full-level training of the GMA model involves two consecutive steps: (1) initial training on age-balanced subsets of the 1 million train/val datasets, utilizing the optimal hyperparameters identified through grid search and (2) subsequent fine-tuning using cell-type balanced data loader on the entire train/val datasets with a reduced learning rate (1/10^th^ of the initial learning rate), building upon the model trained in the first step. During the initial step, a similar training process as the benchmark is applied. In the second step, the pre-trained model from the first step serves as the initial model, and a cell-type balanced data loader is employed to fine-tune the model on the entire train/val dataset. The feature importance calculation and subsequent model applications are based on the refined GMA model obtained after these two stages of training.

### Aging clock models

ScAgeClock comprises five distinct aging clock modules:

‘GatedMultiheadAttentionAgeClock’, ‘MLPAgeClock’, ‘TorchElasticNetAgeClock’,

‘XGBoostAgeClock’, and ‘CatBoostAgeClock’. Each module corresponds to a specific underlying model mentioned above, namely GMA, MLP, Elastic Net, XGBoost, and CatBoost, respectively. These aging clock modules consolidate various functionalities into a single entity, including data loading, model construction, training, validation, testing, and age prediction. Moreover, the GMA, XGBoost, and CatBoost-based aging clock models also provide feature importance calculation functions. Each aging clock model offers a range of tunable parameters to control different aspects of the pipeline, such as data loading, model building, training, and logging information recording. Furthermore, the GMA aging clock model is equipped with checkpoint controls, facilitating large-scale training processes.

### Assessing feature importance and gene expression patterns

Feature importances for boosting-based models are obtained using their native feature importance functions provided by their respective Python packages.

In contrast, feature importances for the GMA aging clock model are derived from the weights of neurons in the feature gating layer. Specifically, the weights are extracted from this layer and their absolute values are calculated for each feature, including embedded features. For categorical features that undergo embedding before passing through the feature gating neural network layer, feature importances are computed as the maximal absolute weights across all corresponding embedded features. In the case of gene features, the absolute weights are used directly as their feature importance values. Additionally, the scAgeClock GMA model also offers alternative methods for calculating categorical feature importances, including taking the mean or sum of the absolute weights from the corresponding embedded features.

To investigate the expression of top-ranked genes by feature importance, 100,000 age-balanced single-cell transcriptome profiles were randomly sampled from the train/val dataset using the H5ADSampler tool from the scAgeClock package.

### Aging Deviation Index

For each cell, the aging deviation index (ADI) is calculated based on the model-predicted age (or transcriptomic-based biological age) and the chronological age of the cell’s donor. The ADI is defined as follows:

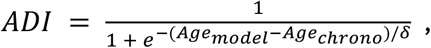

where the 𝐴𝑔𝑒_𝑚𝑜𝑑𝑒𝑙_ represents the model-predicted age, 𝐴𝑔𝑒_𝑐ℎ𝑟𝑜𝑛𝑜_ represents the donor’s chronological age in years, and 𝛿 is a scaling factor with a default value of 25. This calculation can also be performed using the ‘get_ADÌ function in the ‘utility’ tool of scAgeClock.

### Application studies on human brain diseases, normal cell differentiation and cancer cell malignancy

The scAgeClock model’s applicability across diverse biological contexts was investigated by utilizing the scAgeClock ADI to analyze aging acceleration or deceleration patterns in normal human cell types, cells derived from human diseases, and cancer cells representing various tumor phases and metastatic statuses.

The scAgeClock model’s predictions on the testing dataset from CZ CELLxGENE were analyzed to investigate normal human cell type aging dynamics. The ADI was calculated for each cell using the ‘get_ADÌ function from scAgeClock, and the tissue-level cell types were compared based on their ADI scores. Violin plots were generated to illustrate the tissue-level cell types with the highest and lowest average ADIs. For individual-level brain cell type analysis, a normal donor (donor id: H21.33.019) with the largest number of brain cells in the testing dataset was selected. Gene expression data for oligodendrocytes (OLs) and oligodendrocyte precursor cells (OPCs) from this donor were extracted from the testing dataset. Cells with fewer than 200 detected genes and genes expressed in fewer than three cells were excluded for further analysis. The remaining data underwent quality control processing, which included normalization to a total of 1 million UMI counts (Counts Per Million, CPM normalization) followed by log1p transformation. Principal component analysis (PCA) and Uniform Manifold Approximation and Projection (UMAP) analysis were performed using the ‘sc.pp.pcà and ‘sc.tl.umap’ functions from scanpy[31] tool with default settings.

The human disease study was based on human cortex scRNA-seq datasets from the ZEBRA database[22] (more details can be found in the “Data Collection” section). The original cell annotations for cell type (“cell_type2” column in the original metadata) and sex (“sex” column in the original metadata) were matched with scAgeClock’s categorical features “cell_type” and “sex”, respectively. The “age_conti” column from the original metadata was used as the chronological age labels. The data was then formatted using the ‘format_anndata_multiplè function from scAgeClock, which included concatenation of the categorical feature index values into gene expression matrix, gene feature selection, gene expression CPM normalization, and natural logarithm transformation using the log1p function (i.e., log(1+x)). Age predictions were subsequently made using the ‘predict’ function from scAgeClock. Cell-level ADI scores were calculated by the ‘get_ADÌ function from scAgeClock. Disease conditions lacking age information were excluded from the analysis. Additionally, for comparisons of ADI distribution within each age range, only disease conditions with data from at least two donors were included.

To investigate the application of scAgeClock to cancer cells at different tumor phases or with different metastasis statuses, a pan-cancer single-cell RNA sequencing (scRNA-seq) dataset was obtained from a recent human atlas study [23] (data downloading details can be found in the “Data Collection” section). The original data were filtered based on donor age information, and only donors with accurate age information were considered. The columns “platform”, “cellType1”, “tissue” and “sex” were matched with scAgeClock’s categorical features “assay”, “cell_type”, “tissue”, and “sex”, respectively. A similar data formatting process was carried out as for the ZEBRA dataset. Similarly, age predictions were calculated using the ‘predict’ function from scAgeClock. The ADI values were calculated for each cell in the pan-cancer dataset, and ADI values from different tumor phases were compared for each cancer type. To compare the ADI values between cancer cells with or without metastasis, a cancer type with more than 1,000 tumor cells in the metastasis group was selected, which was pancreatic adenocarcinoma (PAAD). The PAAD donor (donorID: PRJCA007744_A032) with the largest number of cells was further selected for individual-level study. To study primary tumor cells with different ADI levels, gene expression data for primary tumor cells (metastasis=no) from this donor were extracted from the whole pan-cancer dataset. Cells with fewer than 200 expressed genes and genes expressed in fewer than three cells were excluded. The data were normalized to CPM as done for the CZ CELLxGENE data and then log1p transformed. The PCA and UMAP analyses were conducted by ‘scanpy.pp.pcà and ‘scanpy.pp.umap’ from scanpy[31] tools, respectively. The ADI mean and standard deviation (SD) of PAAD primary tumor cells were calculated. Primary tumor cells with ADI values less than (ADI mean – ADI SD) were defined as the ADI-lower group, while those with ADI values higher than (ADI mean + ADI SD) were defined as the ADI-higher group. Differentially expressed genes (DEGs) between ADI-higher and ADI-lower primary tumor cell groups were identified by scanpy’s ‘scanpy.tl.rank_genes_groups’ function. P-values were calculated using Wilcoxon rank-sum test and further adjusted by Benjamini-Hochberg (BH) multiple testing correction method. Group-level significant DEGs were defined as those with a BH-adjusted p value less than 0.05 and a fold change larger than 2.

### Functional enrichment analysis

To investigate the functional enrichment of genes that play crucial roles in the scAgeClock model, gene set enrichment analysis (GSEA) was performed using both KEGG (Kyoto Encyclopedia of Genes and Genomes, https://www.genome.jp/kegg/pathway.html) and GO (Gene Ontology, https://geneontology.org) databases. The gseKEGG and gseGO functions from the ClusterProfiler[32] R package were utilized to analyze KEGG and GO enrichments, respectively. The input genes were ranked by their feature importance values in descending order, and the top 3000 genes within the feature importance turning point were further selected for functional enrichment analysis. To visualize gene enrichment on individual pathways, the gseaplot2 function from the ClusterProfiler was employed.

For the ADI-higher and ADI-lower primary cancer cell study, the significant DEGs were defined as those with BH-adjusted P values less than 0.05 and log2 transformed fold changes larger than 1 for each group as described aforementioned. Next, the GO and KEGG pathway enrichment analyses were performed on the up-regulated DEGs for both ADI Group PAAD primary tumor cells using ClusterProfiler’s enrichGO and enrichKEGG function (organism = ’hsa’, pAdjustMethod = ’BH’, and pvalueCutoff = 0.05).

## DATA AVAILABILITY

The human single-cell RNA sequencing (scRNA-seq) datasets used for training, validation, and testing of the scAgeClock model were obtained from CZ CELLxGENE Discover database (census_version=”2024-07-01”, https://cellxgene.cziscience.com). The ZEBRA dataset employed in the brain diseases study were downloaded from the corresponding website (https://ccb-compute.cs.uni-saarland.de/brain_atlas/). Additionally, the human pan-cancer single cell transcriptomic atlas used for cancer cell study was retrieved from Zenodo database (https://zenodo.org/records/15554080).

The scAgeClock model’s related metadata and example datasets are publicly available at https://github.com/gangcai/scageclock. This repository includes example input files in H5AD format, categorical indexing data, model feature data, and pre-trained models, which can be used as a starting point for users to apply the scAgeClock model to their own datasets.

## CODE AVAILABILITY

The source code for the scAgeClock Python package is openly accessible and maintained at https://github.com/gangcai/scageclock. This repository provides related documentation, including installation instructions, basic usage guidelines, and illustrative example use cases to facilitate seamless integration into existing workflows and applications. Furthermore, scAgeClock can be easily installed through via ‘ pip install scageclock’, and is available on PyPI at https://pypi.org/project/scageclock/.

## SUPPLEMENTARY FILES

Supplementary Table S1: Donor metadata summary.

Supplementary Table S2: Cell-type level cell counts.

Supplementary Table S3: Tissue level cell counts.

Supplementary Table S4: Tissue-level cell type performance metrics for the scAgeClock GMA model on testing dataset.

Supplementary Table S5: Ranked feature importances based on scAgeClock GMA model.

Supplementary Table S6: Gene set enrichment analysis of genes ranked by feature importance.

Supplementary Table S7: Differentially expressed genes (DEGs) between ADI-lower and ADI-higher PAAD primary tumor cells.

Supplementary Figure S1: ScAgeClock’s age prediction performance in healthy human donors and the donors with diseases in different brain cell types.

