## Supplementary Figures for "scAgeClock: a single-cell transcriptome based human aging clock model using gated multi-head attention neural networks"

Gangcai Xie^1,2,*^

1. Institute of Reproductive Medicine, Medical School, Nantong University, Nantong, China
2. Jiangsu Province Key Laboratory in University for Inflammation and Molecular Drug Target, Nantong University, Nantong, China

*Correspondence:

Gangcai Xie, Nantong University, Qixiu Road 19, Nantong, China, 226001.

**ABSTRACT**

Aging Clock models have emerged as a crucial tool for measuring biological age, with significant implications for anti-aging interventions and disease risk assessment. However, human aging clock models that offer single-cell resolution and account for cell and tissue heterogeneities remain underdeveloped. This study introduces scAgeClock, a novel gated multi-head attention (GMA) neural network-based single-cell aging clock model. Leveraging a large-scale dataset of over 16 million single-cell transcriptome profiles from more than 40 human tissues and 400 cell types, scAgeClock demonstrates improved age prediction accuracy compared to baseline methods. Notably, the mean absolute error for the best-performing cell type is remarkably low at 2 years. Feature importance analysis reveals enrichment of aging clock genes related to ribosome, translation, defense response, viral life cycle, programmed cell death, and COVID-19 disease. A novel metric, the Aging Deviation Index (ADI) proposed by this study, revealed deceleration of ages in cells with higher differentiation potencies and tumor cells in higher phases or under metastasis, while acceleration of ages was observed in skin cells. Furthermore, scAgeClock is publicly available to facilitate future research and potential implementations.

**Supplementary Figures**

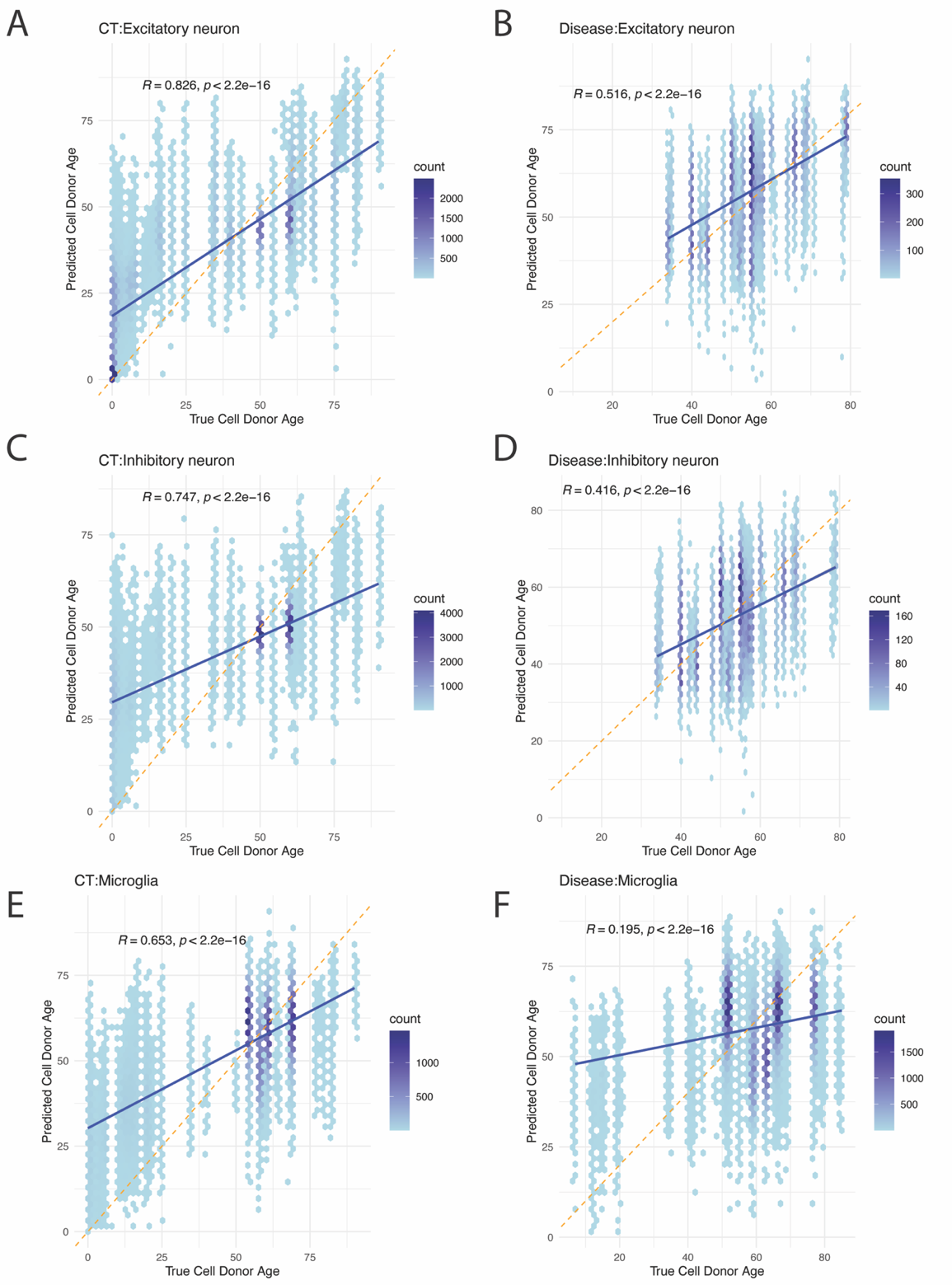

**Supplementary Figure S1: ScAgeClock’s age prediction performance in healthy human donors and the donors with diseases in different brain cell types. (A-F)** Scatterplots illustrating the relationship between chronological ages and predicted ages for human brain excitatory neurons in normal control (A) or disease group (B), for human brain inhibitory neurons in normal control (C) or disease group (D), and for human microglia cells in normal control (E) or disease group (F). Pearson’s correlation coefficient (R) and associated P-value are shown for each plot.

.
